# Electroacupuncture prevents astrocyte atrophy to alleviate depression

**DOI:** 10.1101/2023.02.15.528709

**Authors:** Si-Si Lin, Bin Zhou, Bin-Jie Chen, Ruo-Tian Jiang, Baoman Li, Peter Illes, Alexey Semyanov, Yong Tang, Alexei Verkhratsky

**Author notes:** Send all correspondence to: Si-Si Lin, Yong Tang, Alexei Verkhratsky.

## Abstract

Astrocyte atrophy is the main histopathological hallmark of the major depressive disorder (MDD) in humans and in animal models of depression. Here we show that electroacupuncture prevents astrocyte atrophy in the prefrontal cortex and alleviates depressive-like behaviour in mice subjected to the chronic unpredictable mild stress (CUMS). Treatment of mice with CUMS induced depressive-like phenotypes as confirmed by sucrose preference test, tail suspension test, and forced swim test. These behavioural changes were paralleled with morphological atrophy of astrocytes in the prefrontal cortex, revealed by analysis of 3D reconstructions of confocal Z-stack images of mCherry expressing astrocytes. This morphological atrophy was accompanied with a decrease in expression of cytoskeletal linker Ezrin, associated with formation of astrocytic leaflets, which form astroglial synaptic cradle. Electroacupuncture at the acupoint ST36 as well as treatment with anti-depressant fluoxetine prevented depressive-like behaviours, astrocytic atrophy and down-regulation of astrocytic ezrin. In conclusion, our data further strengthen the notion of a primary role of astrocytic atrophy in depression and reveal astrocytes as cellular target for electroacupuncture in treatment of depressive disorders.

## Introduction

Astrocytes are principal homeostatic cells of the central nervous system (CNS) characterised by a complex spongiform morphology formed by numerous tiny peripheral processes known as leaflets emanating from primary and higher order processes known as branches (Semyanov and Verkhratsky, 2021; Verkhratsky and Nedergaard, 2018). Astrocytic leaflets establish contacts with synapses and form synaptic cradle to foster and maintain synaptic connectivity (Nedergaard and Verkhratsky, 2012; Verkhratsky and Nedergaard, 2014). In particular, leaflets are densely populated by diverse plasmalemmal Na^+^-dependent transporters responsible for neurotransmitter and ion homeostasis in the synaptic cleft (Rose and Verkhratsky, 2016; Verkhratsky and Rose, 2020). In addition, astrocytes secrete numerous factors modulating synaptic function (Augusto-Oliveira et al., 2020). The leaflets demonstrate high degree of physiological morphological plasticity, which also modulates synaptic transmission (Henneberger et al., 2020). In ageing and various forms of neuropathology shrinkage of leaflets leads to reduced homeostatic support, aberrant neurotransmitters spillover, and impaired synaptic plasticity (Henneberger et al., 2020; Popov et al., 2021; Yang et al., 2022). Astrocytic atrophy with reduced leaflets presence is prominent in brain ageing (Popov et al., 2021) and is implicated in pathophysiology of a wide range of neurological diseases (Verkhratsky et al., 2017), including neurodegenerative disorders (Olabarria et al., 2010; Yeh et al., 2011), epilepsy (Plata et al., 2018), addiction (Scofield et al., 2016), and mood disorders (Li et al., 2021). Astrocytic atrophy is the leading histopathological signature of mood disorders in response to stress, in particular in the major depressive disorder (MDD) (Aten et al., 2022; Rajkowska et al., 2013; Rajkowska and Stockmeier, 2013) and post-traumatic stress disorder (Li et al., 2022).

Pharmacological management of MDD remains a challenge, with many patients being resistant to traditional antidepressants. Electroacupuncture (EA) stimulation is employed as an effective and safe therapy to treat multiple diseases including inflammation (Liu et al., 2021; Liu et al., 2020), Parkinson’s disease (Tamtaji et al., 2019), insomnia (Ruan et al., 2009; Yin et al., 2017; Yin et al., 2020), anxiety (Shen et al., 2020) and depression (Jiang et al., 2021; Xu et al., 2020b; Yao et al., 2021). The EA at selective acupoints Dazhui (DU14) and Baihui (DU20) was shown to restore inhibitory/excitatory balance and improve cognition in mouse models of Alzheimer’s disease (Li et al., 2023). Several studies indicated that EA affects astrocytes by modulating expression of glial fibrillary acid protein (GFAP) (Yang et al., 2021) and production of fibroblast growth factor 2 (FGF2), the latter being beneficial for alleviating depressive-like behaviour (Yao et al., 2021).

In the present study, using high-resolution morphological reconstructions we analysed protoplasmic astrocytes in the prefrontal cortex of healthy control mice and mice, which developed depressive-like behaviours following exposure to chronic unpredictable mild stress (CUMS). We found that CUMS lead to a significant morphological atrophy of astrocytes associated with down-regulation of plasmalemmal-cytoskeletal linker Ezrin.

Treatment of these mice with EA or with classical anti-depressant fluoxetin normalised astrocytic morphology, increased expression of Ezrin, and alleviated depressive-like phenotype. Our work suggests that EA may provide a potential non-pharmacological therapeutic tool targeting astrocyte atrophy in MDD.

## Results

### Validation of the depression model: CUMS triggers anhedonia and depressive-like behaviours

Exposure of rodents to CUMS and its variants is widely used to model depression (Aten et al., 2022; Binjie et al., 2022; Liang et al., 2020; Logan et al., 2015). The experimental design including the timeline of the experimental protocol and positions of acupoints used in this study are shown on Fig. 1. Treatment of mice with CUMS regimen for 4 weeks resulted in a significant decrease in sucrose preference reflecting development of anhedonia, one of the key signs of depression in patients (Cooper et al., 2018; De Fruyt et al., 2020; Liu et al., 2018). The sucrose consumption decreased from 0.88 ± 0.01 in control to 0.58 ± 0.05, p < 0.001, n = 6 in mice exposed to CUMS protocol (Fig. 2A). Similarly, exposure to CUMS regimen affected mouse behaviours as revealed in a series of tests. The immobility time in both tail suspension test (TST) and force swimming test (FST) was significantly prolonged (from 100.1 ± 4.8 in control to 155.1 ± 6.3, p < 0.001, n = 6 in CUMS group and from 58.4 ± 4.0 in control to 139.1 ± 3.09, p < 0.001, n = 6 in CUMS group respectively; Fig. 2B,C). In the open field test CUMS decreased exploratory behaviour as demonstrated by a decrease in total running distance, centrepoint cumulative duration and rearing frequency (from 42.488 ± 4.358 m in control to 21.153 ± 1.09 m, p < 0.001, n = 6; from 18.9 ± 1.2 s in control to 6.64 ± 1 s, p < 0.001, n = 6; from 70 ±3.1 in control to 25.5 ± 2.3, p < 0.001, n = 6 respectively; Fig. 2D-G). Together, these data indicate that CUMS induces depressive-like behaviours.

**Figure 1.**
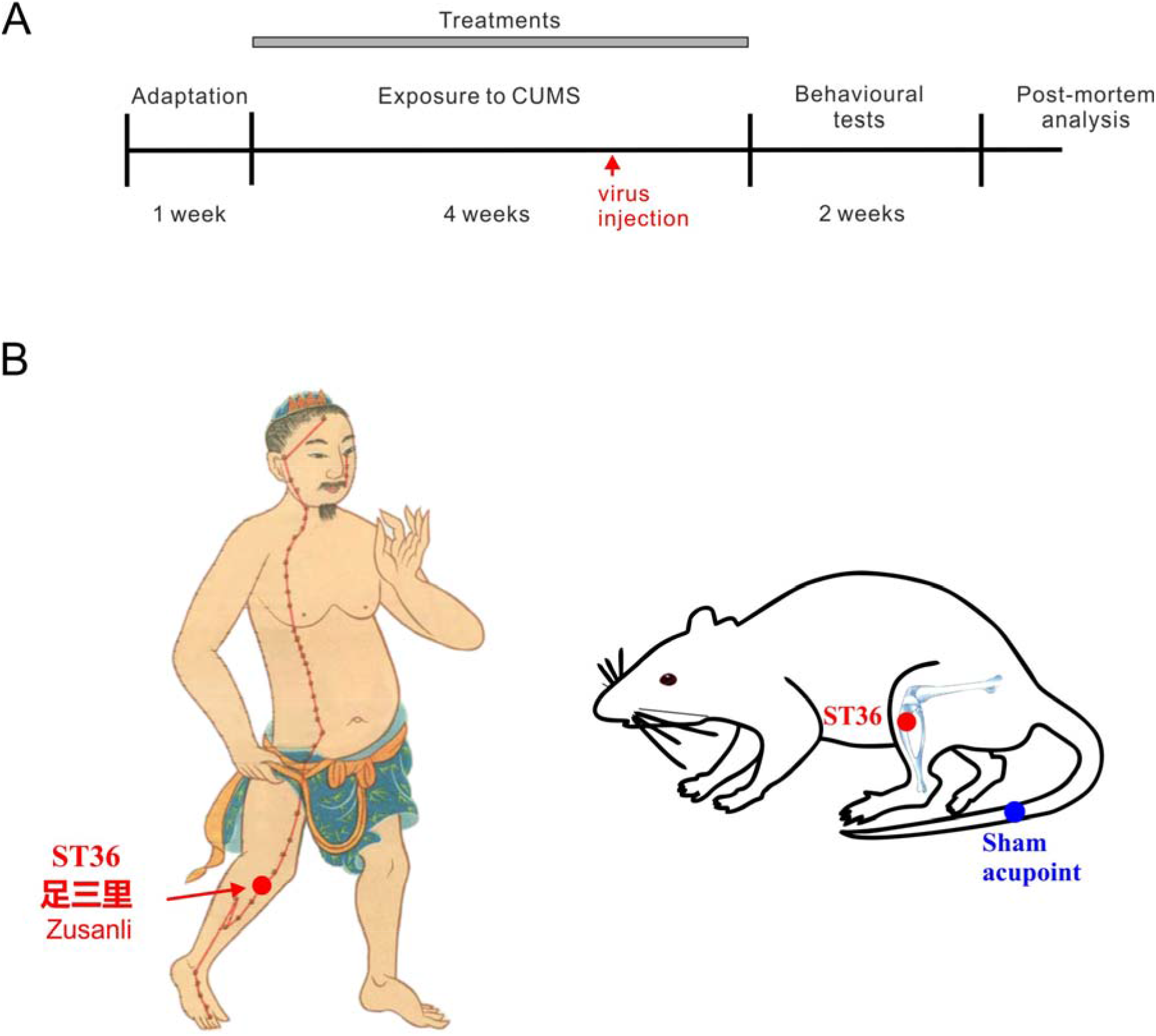
Experimental design. A: Flow diagram of the experimental procedure. B: Location of acupoints used in the study. The man acupoint image is reproduced from an old image created during Ming dynasty; it is reproduced from 新针灸学 *New Acupuncture*, Published by People’s Medical Publishing House in 1954

### Depression is associated with astrocytic atrophy: CUMS reduces morphological profiles and decreases expression of ezrin

To access morphological changes in astrocytes following CUMS we labelled astrocytes in the prefrontal cortex (PFC) of mice by stereotactic injection of AAV5·gfaABC1D·mCherry (Zhou et al., 2019), which allowed reliable morphometric analysis (Fig. 3). The three-dimensional (3D) reconstructions of mCherry-labelled healthy and stressed astrocytes were made from z-stacks of images obtained with confocal microscopy (Fig. 3A). Astrocytic 3D-reconstructions were subjected to Scholl analysis inbuilt in the Imaris software. Exposure to CUMS regimen resulted in a substantial decrease in astrocytic size and complexity (Fig. 3). The maximal number of intersections was significantly decreased form 13.5 ± 0.9 in healthy to 8.2 ± 0.4 in depressed animals (p < 0.001; n = 15, Fig 3B, C), while and the length of astrocytic branches decreased from 11.6 ± 0.7 μm in control to 8.0 ± 0.2 μm, p < 0.001in CUMS mice; n = 15 (Fig. 3D). Similarly, exposure to CUMS protocol significantly reduced the size of astrocytic territorial domain (from 1151.6 ± 44.3 μm^2^ in control to 283.7 ± 13.3 μm^2^ in CUMS animals, p < 0.001; n = 6, Fig 3E,F).

Ezrin, a member of ezrin-radixin-moesin (ERM) family tethers actin filaments to the plasma membrane, and is essential for formation and structural plasticity of astrocytic leaflets and astrocyte-synaptic interactions (Badia-Soteras et al., 2022; Derouiche and Geiger, 2019). We quantified ezrin expression with immunocytochemistry, applied to the PFC preparations from mice subjected to mCherry astrocytic labelling (Fig. 4 A). The fluorescence of ezrin associated with mCherry-positive astrocytic profiles was significantly decreased in depressed animals (from 1201.2 ± 73.4 in control to 813.0 ± 28.4 in CUMS-treated animals, p < 0.01; n =15, Fig 4B, C). The number and surface area of ezrin-positive puncta uniformly decreased around the soma and branches of depressed animals (from 94.7 ± 16.3 μm^2^ to 51.0 ± 6.8 μm^2^, p = 0.016, n = 15, and from 204.8 ± 36.6 μm^2^ to 110.2 ± 13.5 μm^2^, p = 0.018, n = 15 respectively, Fig. 4D, E).

### Fluoxetine and EA prevent depressive-like behaviours induced by CUMS

In this study we compared the anti-depressant efficacy of EA in the specific acupoint with the action of classic anti-depressant drug fluoxetine (a.k.a Prozak). We found that treatment with fluoxetine as well as treatment with EA at Zusanli acupoint 足三里 ST36) used in traditional Chinese medicine (TCM) for therapy of depression fully prevented the development of depressive-like behaviours in CUMS exposed mice (Fig. 5). In particular, in animals treated with fluoxetine and EA sucrose consumption was significantly higher compared to the CUMS group (Fig 5A; fluoxetine: 0.8 ± 0.03 vs. 0.6 ± 0.05 in CUMS, p < 0.001 EA: 0.8±0.03 vs. 0.6±0.05 in CUMS; p < 0.01, n = 6). Similarly, fluoxetine and EA prevented the development of depression-like behaviours in TST and FST (Fig 5B, C); both treatments also prevented a decrease in exploratory behaviours in the open field test (Fig. 5D-G). The efficacy of fluoxetine and EA were similar without any significant defences in tests readouts. At the same time, neither injection of PBS nor sham electroacupuncture (EA1, EA2: acupuncture without electrical stimulation or acupuncture in clinically irrelevant acupoints, see methods section) were effective in preventing the development of depressive-like behaviours in mice exposed to the CUMS protocol.

### Fluoxetine and EA prevent astrocyte atrophy and increase the presence of ezrin

In parallel with preventing development of depression-like behaviours, treatment with fluoxetine and exposure to EA averted CUMS-induced astrocytic atrophy (Fig. 6) and CUMS-induced decrease of astrocyte-associated ezrin (Fig. 7). In particular, in the animals subjected to fluoxetin or EA treatments maximal number of intersections and length of astrocytic branches was significantly higher than in the CUMS group (Intersections; fluoxetine: 14.2 ± 0.5 vs. 8.2 ± 0.4 in CUMS; p < 001, n = 15; EA: 17 ± 1.0 vs. 8.2 ± 0.4 in CUMS, p < 0.001, n = 15; length of branches fluoxetine: 13.6 ± 0.6 vs. 8.0 ± 0.2 in CUMS; p <0.001, n = 15; EA: 14.0±0.7 vs 8.0±0.2 to p < 0.001; n = 15; Fig. 6A-C). Likewise, both fluoxetine and EA prevented CUMS-induced decrease in astrocytic territorial domain (fluoxetine: 790.5 ± 43.2 vs. 283.7 ± 13.4 in CUMS; p < 0.001, n = 6; EA: 1078.0 ± 62.3 μm^2^ vs. 283.7 ± 13.4 μm^2^ in CUMS; n = 6; Fig. 6D). Neither injections of PBS, nor sham EA were able to prevent CUMS-induced changes in astrocytic morphology (Fig. 6B-D).

Similar effects fluoxetin and EA exerted on ezrin association with astrocytes (Fig. 7). In animals receiving fluoxetin or EA treatment the fluorescence intensity of ezrin associated with astrocytic domain was significantly higher than in CUMS only group (fluoxetine: 1248.4 ± 140.5 vs. 703.8 ± 19.4 in CUMS; p < 0.01 n = 15; EA: 1176.14 ± 254.6 vs. 703.8 ± 19.4 in CUMS, p < 0.05; n = 15, Fig. 7B). Immunolabelled ezrin puncta increased homogeneously in soma and branches areas (Fig. 7B). Again, neither injections of PBS, nor sham EA affected ezrin levels.

## Discussion

### Chronic stress induces astrocytic atrophy: pathophysiological relevance for depression

Using high-resolution morphometric analysis we confirmed pervious observations showing that chronic stress causes morphological atrophy of astrocytes in several areas of the brain including PFC (Aten et al., 2022; Codeluppi et al., 2021; Tynan et al., 2013), which was at the focus of our study. The high-resolution morphometry of astrocytes *in situ* requires cytosolic labelling with fluorescent probe; the latter could be either injected into the cells of interest through microelectrode (for example Lucifer yellow (Zhou et al., 2019)), or patch-pipette (for example Alexa Fluor 594 (Popov et al., 2020)) or expressed under the control of astroglia-specific promoter (such as *Aldh111* (Aten et al., 2022) or *Gfap* (Codeluppi et al., 2021)). In our study we used mCherry and *Gfap* promoter for a viral transfection (AAV5-gfaABC1D-mCherry; see also (Zhou et al., 2019)). Astrocytic atrophy was quantified by Scholl analysis, which showed significant decrease in the number of intersections, by decreased length of astrocytic branches and by reduced area of astrocytic territorial domains (Fig. 2). In addition, chronic stress affected astrocytic presence of ezrin, as judged by immunocytochemistry – in animals exposed to CUMS number of labelled ezrin puncta associated with individual astrocytes (both at the soma and at branches) was significantly decreased (Fig. 3). Ezrin is a linker of plasmalemma and cytoskeleton, and it is critical for formation and morphological plasticity of astrocytic leaflets (Derouiche and Geiger, 2019; Schacke et al., 2022); decrease in ezrin signals decrease in the number and/or volume of leaflets. The leaflets are part of idiosyncratic morphology of protoplasmic astrocytes, which gives them spongiform or bushy appearance (Semyanov and Verkhratsky, 2021). These leaflets are characterised by an extremely high surface-to-volume ratio and absence of organelles (Gavrilov et al., 2018; Semyanov and Verkhratsky, 2021), which stipulates specificity of ionic signalling in these compartments (predominance of Ca^2+^ entry and prominence of Na^+^ signalling (Lim et al., 2021; Verkhratsky et al., 2020)). The leaflets are closely associated with synapses and, by virtue of high density of plasmalemmal homeostatic transporters, sustain and regulate synaptic transmission (Verkhratsky and Nedergaard, 2018).

**Figure 2.**
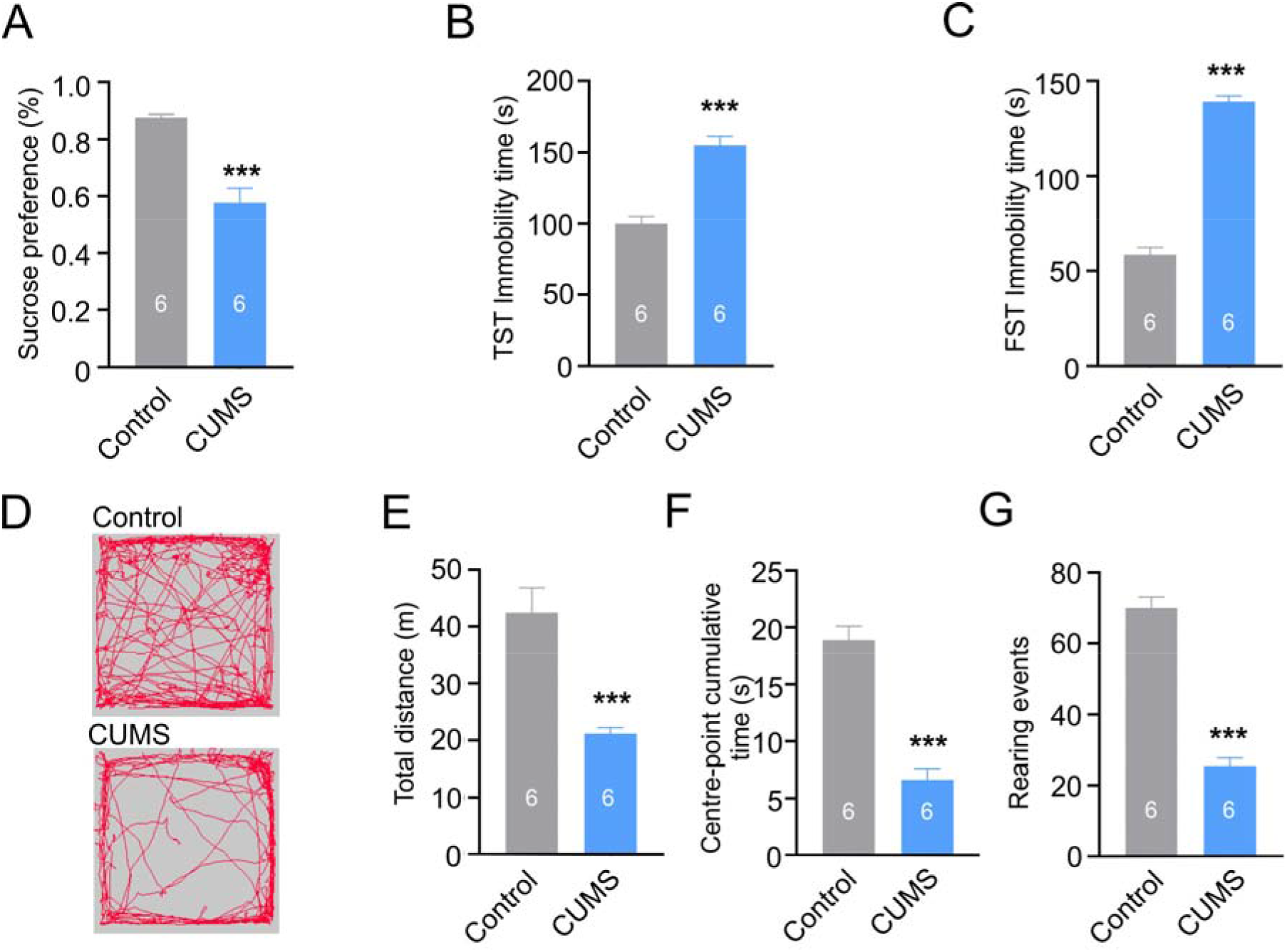
The CUMS regimen triggers depressive-like phenotypes in mice. A-C: CUMS reduced sucrose intake and prolonged the TST and FST immobility time. E - F: CUMS affected exploratory behaviour in an open field test; E: representative running trace in open field test, the observation time was 10 min. F-H: total running distance, centre-point cumulative duration, and number of rearing events. All data is presented as mean ± s.e.m. * p<0.05, ** p<0.01, *** p<0.001. The number of experiments is indicated on each column.

Astroglial atrophy, asthenia and loss of function is a distinct astrogliopathological entity observed in many neurological diseases (Verkhratsky et al., 2017). Decreased astrocytic presence in the brain active milieu has multiple consequences. Shrunken astrocytes open diffusional channels and facilitate volume transmission (Verkhratsky et al., 2021); at the level of single synapses, reduced presence of astrocytic leaflets impairs clearance of glutamate, thus leading to increased availability (with possible excitotoxic repercussions) and spillover of the latter thus affecting synaptic plasticity (Popov et al., 2021). Furthermore, reduced glial presence limits homeostatic support, provision of glutamine, K^+^ buffering, control over pH and supply of scavengers of reactive oxygen species (Verkhratsky et al., 2019; Verkhratsky and Nedergaard, 2018). To the contrary, when the growth of astrocytic leaflets is promoted, the support of the synaptic transmission is increased. Astrocytes are characterised by high degree of morphological plasticity and various environmental factors are known to increase astrocytic domains, mainly through an increase of leaflets presence. These environmental factors include, for example enriched environment and physical activity (Carvalho-Paulo et al., 2017; Rodriguez et al., 2013), dieting (Popov et al., 2022; Popov et al., 2020) and other life style factors, positively impacting cognitive abilities (Augusto-Oliveira and Verkhratsky, 2021). In the context of mood disorders astrocytic atrophy can contribute to the aberrant neurotransmission and affect neuronal survival and overall functional performance. Indeed, specific manipulation with astrocytes and astrocytic homeostatic cascades can induce depression-like behaviours, whereas treatment with antidepressants rescues these behaviours and reverses astrocytic atrophy (Banasr and Duman, 2008; John et al., 2015). Here we demonstrate that EA acts very similarly to chemical antodepressants and prevents development of depressive behaviours and astrocytic atrophy in the PFC of mice exposed to CUMS.

### Electroacupuncture acts on astrocytes to prevent depression?

Acupuncture was used in Traditional Chinese Medicine as a therapeutic manipulation for last 3000 to 4000 years (Ma et al., 2021); with the first codex formalising acupuncture treatments (*The Yellow Emperor’s Classic of Internal Medicine*) was published around 100 BC (Ke, 2022). Acupuncture is now used worldwide for treatment of various medical conditions (Jang et al., 2020; Liu et al., 2021; Tang et al., 2016; Xu et al., 2020a). Acupuncture is employed for therapies of neuropsychiatric diseases, including depression (Jung et al., 2021; MacPherson et al., 2013) and anxiety (Amorim et al., 2022). In particular, 6 weeks of acupuncture at acupoints Baihui (GV20) and Zusanli (ST36), or Taichong (LR3), Sanyinjiao (SP6), Neiguan (PC6), and Shenmen (HT7), was as effective as oral fluoxetine in treating depression, although acupuncture showed better response and improvement rates (Sun et al., 2013). Similarly, acupuncture at ST36 and CV4 acupoints was effective in alleviating depressive-like behaviours in animal models (Eshkevari et al., 2015; Le et al., 2016). In our present study we chose to perform EA at the Zusanli (ST36, 足三里, or point of longevity) acupoint, known to be linked to the brain. Acupuncture at this acupoint affects brain metabolism of glucose (Yin et al., 2003), modulates excitation-inhibition balance in cortex (Sun et al., 2019) and triggers responses of several brain areas including the anterior cingulate cortex (ACC), ventrolateral prefrontal cortex (VLPFC), occipital cortices, somatosensory cortex, and midbrain (Hui et al., 2005; Liu et al., 2010).

We found that EA, as well as treatment with fluoxetine fully prevents the development of depressive-like behaviours in mice subjected to chronic stress (Figs 2, 5). To elucidate the possible brain cellular target of the EA, we performed in depth analysis of fine morphology of cortical astrocytes, known to become atrophic in depression. Exposure of mice to the CUMS regimen led, as expected, to a significant decrease in astrocytic size and complexity (Fig. 3), which were fully prevented by both fluoxetine treatment and EA at acupoint ST36. This effect was specific as sham acupuncture was ineffective. Treatment with EA (as well as with fluoxetine) prevented decrease in ezrin associated with astrocytic structires. We may therefore suggest that chronic stress leads to a retraction of astrocytic leaflets and decrease of astrocyte-synaptic association, which in turn impairs synaptic transmission and plasticity, as has been demonstrated in ageing and various neuropathologies (Verkhratsky et al., 2019, 2021, Plata et al., 2018, Popov et al., 2021) responsible for generation of depressive-like behaviours. Our data are supported by recent observation demonstrating that EA restores expression of astrocyte-specific glutamate transporter EAAT2 following chronic stress; this restoration developed in parallel with amelioration of depressive-like behaviours (Luo et al., 2017; Tu et al., 2019).

**Figure 3.**
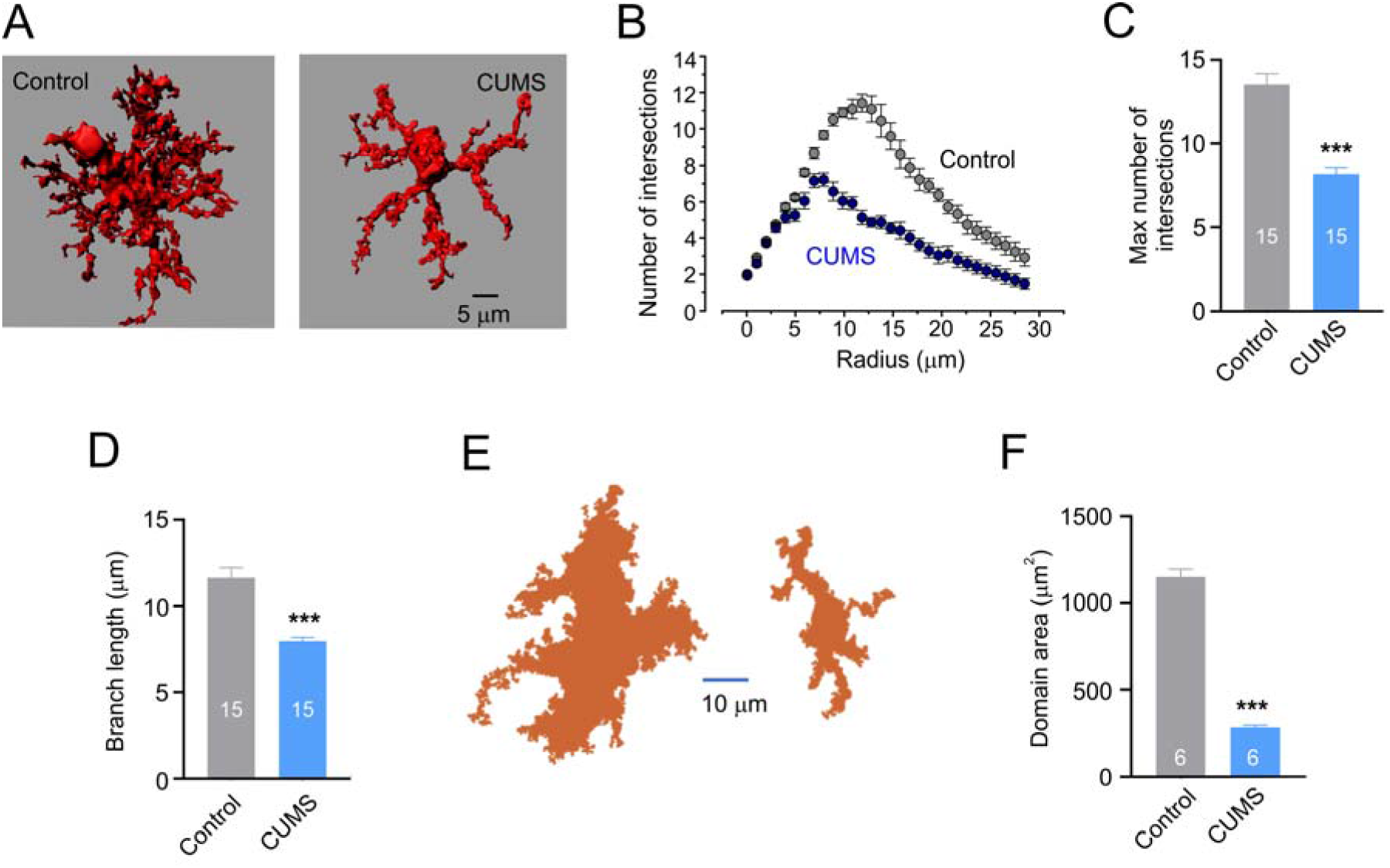
CUMS induces morphological atrophy in PFC astrocytes. A: Representative 3D reconstruction of astrocyte in control and CUMS groups. B: Sholl analysis of astrocytic morphology for control and CUMS groups shows the number of intersections of astrocytic branches with concentric spheres centred in the middle of cell soma. C: Maximal number of intersections for astrocytes in control and CUMS groups. D: Average length of astrocytic processes in control and CUMS groups. B-D: n = 15 for each group. D: Representative examples of astrocytic territorial domains obtained as a projection of astrocytes along the z-axis projection for control and CUMS animals. E,F: Average astrocytic domain area (E) and average length of astrocytic processes for control and CUMS group. All data is presented as mean ± s.e.m. * p<0.05, ** p<0.01, *** p<0.001. The number of experiments is indicated on each column.

**Figure 4.**
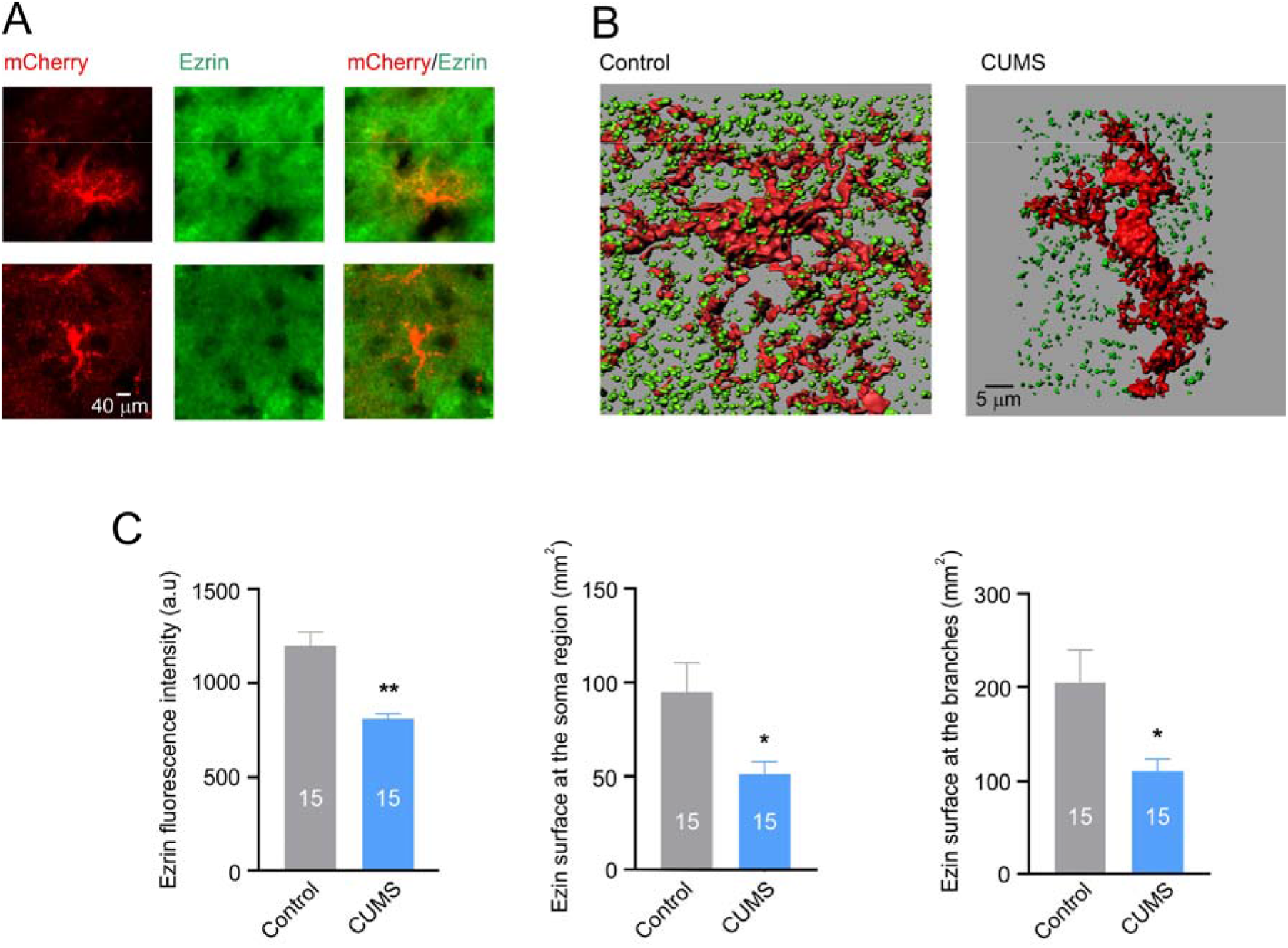
Chronic stress decreases ezrin association with astrocytes. A: Images of astrocytes labelled with AAV-GfaABC1D-mCherry and immunostained for ezrin. B: Representative 3D-reconstruction of astrocytic profiles (red) with Ezrin puncta (green) for control and CUMS groups. C: Average fluorescence intensity of ezrin and surface area of ezrin puncta associated with soma and branches of 3-D reconstructed astrocytes. All data is presented as mean ± s.e.m. * p<0.05, ** p<0.01, *** p<0.001 compared to control group. The number of experiments is indicated on each column.

What are the mechanisms translating acupuncture into changes in astrocytic morphology and functional boost which protects against depressive changes instigated by chronic stress? At this stage we can only speculate. As we demonstrated in this study, classic anti-depressant fluoxetine rescues stress-induced depressive behaviours as well as prevents astrocytic atrophy. Fluoxetine acts on astrocytes either through inhibition of serotonin transporter SERT/ SLC6A4 (which, however, is mainly expressed in neurones) or through acting directly on astrocytic 5-HT_2B_ receptors. Activation of these receptors by fluoxetine triggers several signalling cascades (Hertz et al., 2015; Hertz et al., 2014), and induces transactivation of epidermal growth factor receptors (EGFR), which, in turn, recruits MAPK/ERK or PI3K/AKT downstream cascades to regulate expression of several genes (such as Ca^2+^-dependent phospholipase A2, cPLA2, subtype 2 of adenosine deaminases acting on RNA’s, ADAR2, or subtype 2 of kainate receptors, GluK2) related to mood disorders (Peng et al., 2015). Fluoxetine also normalises interstitial pH which is affected in mood disorders by phosphorylating and stimulating astrocytic Na^+^-H^+^ transporter NHE/SLC9a1 (Ren et al., 2015). Whether EA may act through the similar cascade requires further investigation.

### Conclusion

In summary, we demonstrated that EA prevents the development of depression-like behaviours in mice exposed to chronic stress. At cellular and molecular levels, EA prevents morphological atrophy of astrocytes (which is a leading histopathological hallmark of depression) as well as the decrease in astrocyte-associated ezrin that controls fine morphology of these cells. Our study therefore provides, for the first time, experimental evidence for an idea that EA boosts astrocytic presence in the brain active milieu thus protecting it against stress-induced pathological deformations resulting in depressive behaviours.

## Materials and methods

### Animals

All experiments were performed on C57BL/6 mice (obtained from Chengdu Dossy Experimental Animal Co., Chengdu, China); mice were 7 weeks old at the beginning of the experimental protocol, which lasted 7 weeks (Fig. 1A). All mice were adapted to the standard laboratory conditions (24 ± 2°C room temperature and 65 ± 5% humidity on 12/12 h light-dark cycles) with drinking water and food available ad libitum. The experimental procedures were made in accordance with the National Institute of Health Guidelines for the Care and Use of Laboratory Animals and approved by the Animal Ethics Committee of Chengdu University of Traditional Chinese Medicine(protocol code, AM3520, 8 May 2019).

### Chronic unpredictable mild stress (CUMS) regimen

Mice were exposed to the random sequence of stressors during each 24-h period for 4 weeks, as previously described (Dong et al., 2015; Liang et al., 2020). These stressors included water and food deprivation (12 h), cage tilt 45° (12 h), group housing (12 h), swimming in 4 °C water (5 min), foot shock (1mA, 5 min), noise (120 dB for 3 h), tail suspension (5 min), damp bedding (12 h), cage shaking (40/min for 5 min), and restraint (1h).

### Experimental groups and treatments

We established 8 experimental animal groups: (i) control group; which did not receive any interventions; (ii) CUMS groups – animals exposed to CUMS only; (iii) CUMS + fluoxetine group – animals exposed to CUMS and weekly injections (for all period of CUMS treatment) of fluoxetine at 10 mg/kg; (iv) CUMS + PBS group, animals exposed to CUMS and daily injections of PBS (0.2 ml); (v) EA group animals exposed to EA daily and (vi-vii) sham EA1 and EA2 groups exposed to sham acupuncture daily.

EA was administered at roughly the same time of the day (10:00 a.m. to 11:00 a.m.) to awake animals, immobilised by two Velcro brand hooks and loop fasteners as well as additional tapes fixed to a wooden block for the duration of EA (Zhang et al., 2020). EA was delivered to the therapeutically relevant Zusanli acupoint (ST36; located at the knee, about 2 mm for mice below the fibular head). An electrical current of 0.5 mA and a frequency of 2 Hz was delivered for 30 min, by an acupoint nerve electrostimulator (HANS-200, Nanjing Jisheng Medical Technology Co., Jiangsu, China). EA was applied through stainless steel needles (2.5 cm long, 0.25 mm diameter; Hwato-Med. Co., Jiangsu, China), introduced 2–3 mm deep below the skin at ST36 unilaterally. Sham treatments were as follows. In the EA1 sham groups the needle was inserted but the electrical stimulation was not applied. In the EA2 sham groups the needle was positions at the nonacupoint at the tail (Torres-Rosas et al., 2014).

### Behavioural tests

#### Sucrose preference test

The sucrose preference test is a reward-based test and a measure of anhedonia, as previously described (Zhao et al., 2022). The mice were singly caged for 3 days and given two 50 mL bottles containing water or water-based 1% sucrose solution (wt/vol), respectively. The bottle positions were switched daily to avoid a side bias. Following a 24 h period of water and food deprivation, the preference for sucrose or water was determined overnight. Sucrose preference (%) was quantified as (vol sucrose/(vol sucrose + vol water)) × 100%.

#### Tail Suspension Test

The tail suspension test is a behavioural despair-based test assessing the duration of immobility of mice subjected to inexorable conditions, as previously described (Zhao et al., 2017). Each mouse was suspended by its tail at a height of 20-25 cm by using a piece of adhesive tape wrapped around the tail 1 cm from the tip. Behaviour was recorded for 6 min. The duration of immobility was calculated by an observer blinded to the treatment groups. The mice were considered to be immobile only when they remained completely motionless; mice that climbed along their tails were not included.

#### Forced Swimming Test

The FST was performed as previously described (Zhao et al., 2022), in a clear glass cylinder filled with water (temperature, 23 - 25°C); cylinder’s dimensions were: height, 30 cm; diameter, 20 cm; water level, 15 cm. Mice were gently placed in the tanks. The duration of immobility within the 6 min of observation was determined. The movement of the animals was video recorded and analysed later. Following the swimming session, the mice were removed from the water by their tails, gently dried with towels, and kept warm under a lamp in their home cages. They were considered to be immobile whenever they stopped swimming and remained floating passively, still keeping their heads above the surface of the water.

#### Open field test

The open field test was performed as previously described (Zhao et al., 2022). The apparatus consisted of a rectangular chamber (50 × 50 × 50 cm) made of white, high density, non-porous plastic. Mice were gently placed in the centre of the chamber and their motility was recorded for 10 min. The total running distance, and the time spent in the centre versus the periphery of the open field chamber were recorded by a camera connected to a computer using an automated video tracking program (EthoVision XT 9.0; Noldus, Wageningen, The Netherlands). The chamber was thoroughly cleaned with 95% ethanol, and dried prior to use and before subsequent tests, to remove any scent clues left by the previous subject.

### AAVs microinjections

Viral injections were performed at the end of week 3 of CUMS treatment as indicted in Fig. 1 by using a stereotaxic apparatus (RWD, Shenzhen, China) to guide the placement of a Hamilton syringe fixed with bevelled glass pipettes (Sutter Instrument, 1.0-mm outer diameter) into the PFC (Soiza-Reilly et al., 2019). The injection site was located at half of the distance along a line defined between each eye and the lambda intersection of the skull. The needle was held perpendicular to the skull surface during insertion to a depth of approximately 0.2 mm. A total of 0.7 μl of AAV5-gfaABC1D-mCherry (1× 10^12^ gc/mL; Taitool Bioscience, Shanghai, China), was slowly injected into right sides of the PFC. Glass pipettes were left in place for at least 5 min. After injection, animals were allowed to completely recover under a warming blanket and then returned to the home cage.

### Immunohistochemistry

Mice were perfused in the morning with cold paraformaldehyde (PFA, 4% w/v in phosphate buffer saline (PBS)) under deep isoflurane (2%, 5 min) and pentobarbitone (1%, 50 mg/kg) anaesthesia. Brains were collected, postfixed and cryoprotected in 30% (w/v) sucrose solution. Brains were cut using a cryostat in 45 um thick sections; slices were immediately transferred into storing solution (30% w/v sucrose and 30% ethylene glycol in PBS) and kept at 80 °C until use. Free floating sections were incubated 1 h in saturation solution (6% foetal calf serum in PBS). The sections were then incubated overnight in the same solution complemented with the primary antibody (rabbit anti-Ezrin 1:100 CellSignalling, Danvers, Massachusetts, USA). After washing in PBS three times, slices were incubated 1 h at 37 □ in saturation solution containing the relevant secondary antibody (goat anti-rabbit Alexa 488; Invitrogen, Carlsbad, California, USA). After washing in PBS 3 three times, labelling nucleus with DAPI, the coverslips were mounted on slides using anti-fade solution (Solarbio, Beijing, China). Confocal microscopy (Olympus, Tokyo, Japan) or normal fluorescence microscope (Leica, Wetzlar, Germany) were used to obtain images.

### Sholl analysis

Sholl analysis is a commonly used methods to quantify astrocyte processes complexity (Codeluppi et al., 2021; Popov et al., 2022). All processing steps were performed using image analysis software ImageJ [https://imagej.net/imagej-wiki-static/Sholl_Analysis]. In brief, Z-stacks corresponding to the emission spectrum (565–610 nm) of mCherry-labelling (resolution was 512 × 512 pixels (0.2 μm/px) on XY axis with a step on Z-axis 1 μm/frame were re-sampled to the same lateral resolution of 0.25 μm/px.

### 3D reconstructions

The confocal imaging stacks were collected with a Z-step size of 0.25 μm under a confocal microscope (Olympus, Tokyo, Japan). Three-dimensional reconstructions were processed offline using Imaris 7.4.2 (Bitplane, South Windsor, CT) as reported previously (Octeau et al., 2018). In brief, the astrocyte soma and processes were measured and reconstructed according to its own parameter. Processes diameter was measured as one-tenth of astrocyte soma. In addition, Ezrin was measured as 1 mm in every group. The surface-surface colocalisation was calculated by a specific plugin of Imaris Zhou, 2019 #37}.

### Statistics

All statistical analyses were performed by Graphpad prism 8. All data were expressed as means ± SEM of n observations, where n means the number of animals in behavioural tests, or astrocyte cell from at least three animals. Data with more than two groups were tested for significance using one-way ANOVA test followed by the Holm–Sidak test. Multiple comparisons between the data were performed in case of their non-normal distribution, using the Kruskal–Wallis ANOVA on ranks, followed by the Tukey’s test. A two-way ANOVA followed by the Dunn’s test was performed to compare data obtained in Figure 2I, Figure 3B, Figure 6B. Significance was defined as P < 0.05.

**Figure 5.**
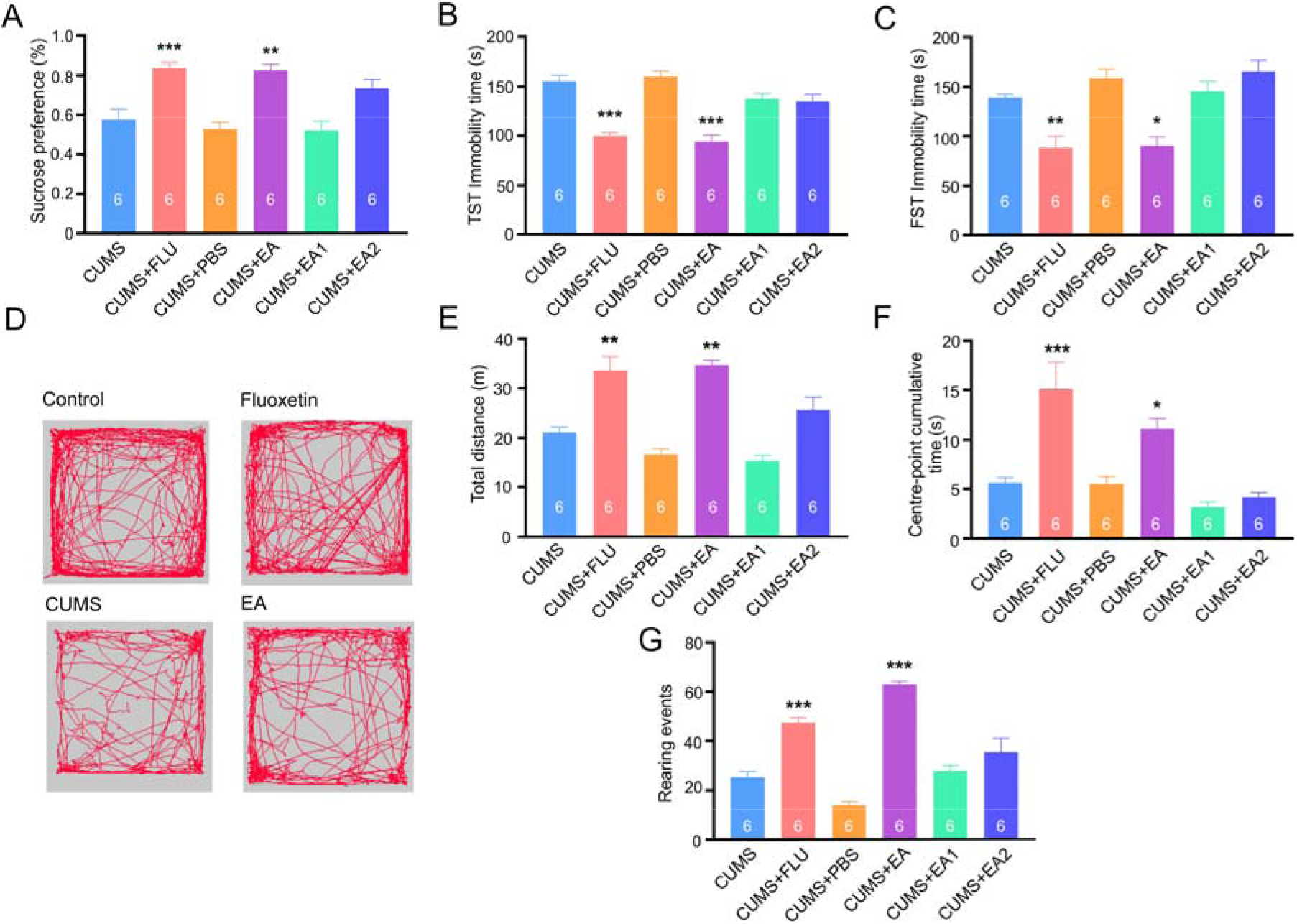
Treatments with fluoxetine and EA prevents chronic stress-induced development of depressive-like behaviours. A: Sucrose preference test. B: TST immobility time. C: FST immobility time. D: Representative running trace in open field test, the observation time was 10 min. E-G: Average values for total distance, centre-point time and number of rearing events. Experimental groups are indicated on the graphs. All data is presented as mean ± s.e.m. * p<0.05, ** p<0.01, *** p<0.001 compared to control group. The number of experiments is indicated on each column.

**Figure 6.**
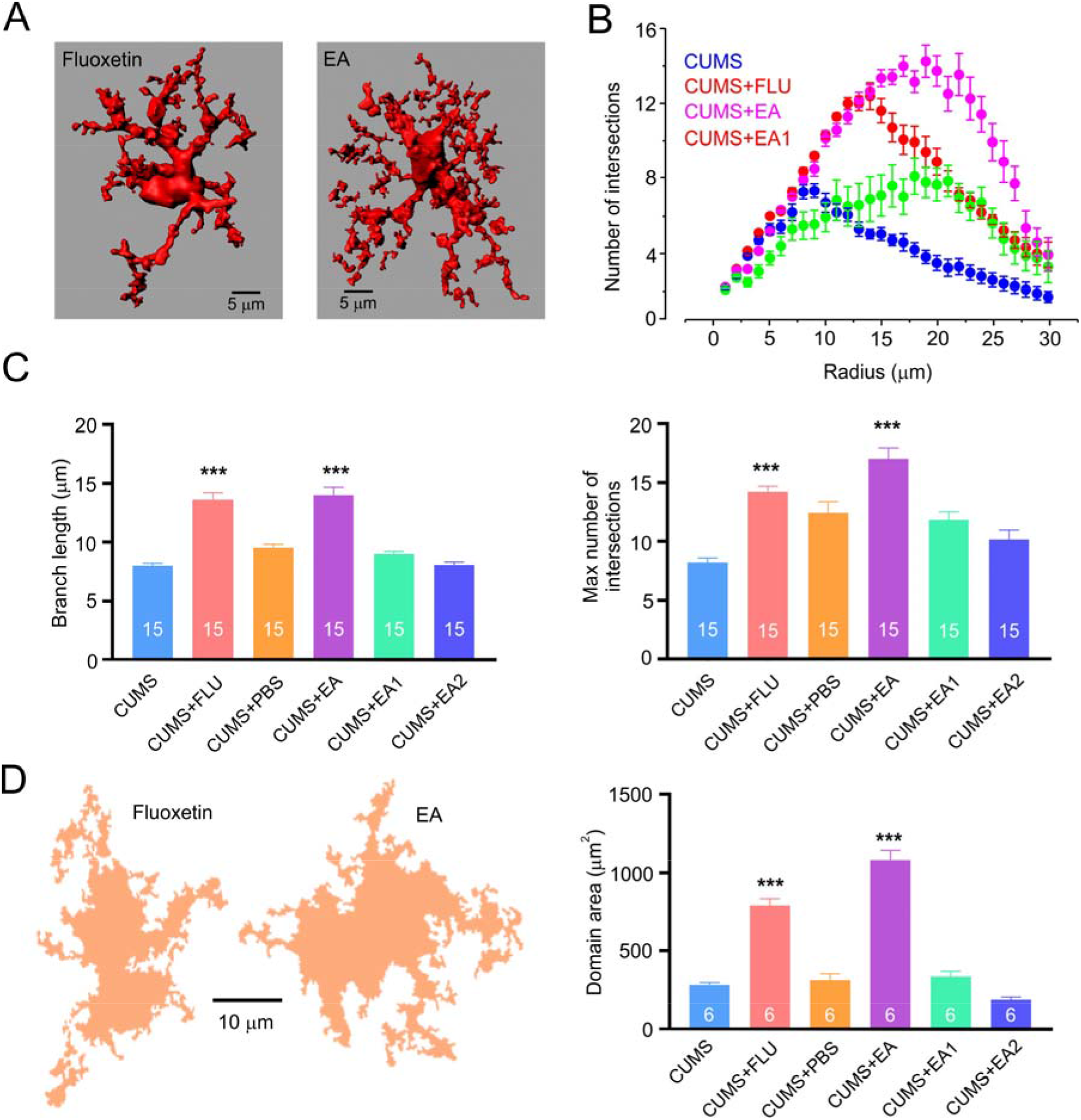
Treatment with fluoxetine and EA prevents chronic stress-induced development of astrocytic atrophy. A: Representative 3D reconstruction of astrocyte in CUMS+fluoxetine and CUMS+EA groups. B: Sholl analysis of astrocytic morphology for CUMS, CUMS+fluoxetine, CUMS+EA and CUMS+EA1 groups.. C: Maximal branch length and number of intersections; experimental groups are indicated on the graph. B, C: n = 15 for each group. D: Representative examples of astrocytic territorial domains obtained as a projection of astrocytes along the z-axis projection for CUMS+fluoxetine and CUMS+EA. The average domain area across various experimental groups is shown on the left. Experimental groups are indicated on the graphs. All data is presented as mean ± s.e.m. * p<0.05, ** p<0.01, *** p<0.001 compared to control group. The number of experiments is indicated on each column.

**Figure 7.**
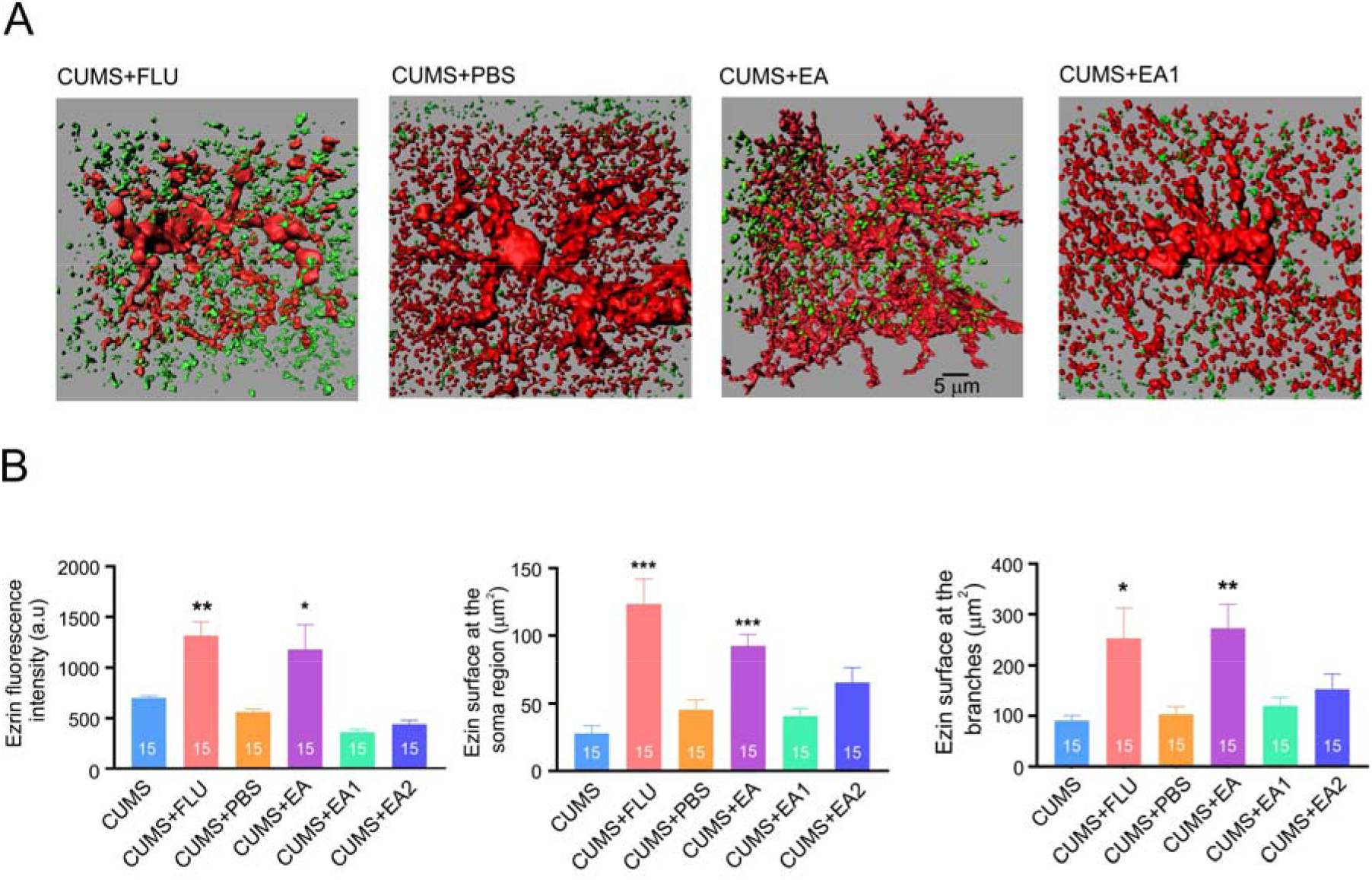
Treatment with fluoxetine and EA preserves ezrin-astrocyte association under CUMS regimen. A: Representative 3D-reconstruction of astrocytic profiles (red) with Ezrin puncta (green) for CUMS+fluoxetine, CUMS+PBS, CUMS +EA and CUMS+EA1 (sham acupuncture procedure). B: Average fluorescence intensity of ezrin and surface area of ezrin puncta associated with soma and branches of 3-D reconstructed astrocytes across various experimental groups. Experimental groups are indicated on the graphs. All data is presented as mean ± s.e.m. * p<0.05, ** p<0.01, *** p<0.001 compared to control group. The number of experiments is indicated on each column.

## Acknowledgments

This work was supported by grants from NSFC-RSF (82261138557), NSFC (82274668, 82230127), the Innovation Team and Talents Cultivation Program of the National Administration of Traditional Chinese Medicine (ZYYCXTD-D-202003), and the Sichuan Science and Technology Program (2022YFH0006).

## Author contributions

AV, YT, and SL conceived the study, AV and SL designed experiments, SL performed experiments, BL and BJC provided expertise in CUMS, BZ and RTJ contributed to confocal experiments and image analysis, AS and PI contribute to discussion, AV and SL wrote the manuscript, AV, YT, PI, AS, BL edited the manuscript.

